# Effect of Intermittent Pneumatic Compression on Preventing Deep Vein Thrombosis Using Microfluidic Vein Chip

**DOI:** 10.1101/2023.04.12.536669

**Authors:** Hongtao Dai, Senlin Chai, Wei Xu, Yao Yao, Wenlai Tang, Jianping Shi, Ji quan Yang, Qing Jiang, Liya Zhu

## Abstract

Deep vein thrombosis (DVT) is a common disease often occurring in the lower limb veins of bedridden patients. Intermittent pneumatic compression (IPC) has been considered an effective approach to solve this problem. In our previous research, 264 patients were randomly treated either with IPC for one or eight hours per day. The incidence of severe venous thrombosis was significantly lowered in the IPC group than in the control group. However, it is still a chanllenge to real-time monitoring the blood flow and thrombus formation process during IPC operation periods. Here we made a microfluidic vein chip with valves embedded in a flexible channel that can mimic the compression of vessels by IPC contractrion. Driven by inlet blood velocity obtained clinally, numerical simulation were conducted to identify the shear stress and laminar pressure distribution in the vein. The obtained results showed that the thrombus formation can be reduced with Higher compressive pressure and smaller time interval. 24s interval time and 40mmHg maximum contractive pressure was considered to be the most appropriate parameters for DVT prevention. This vein chip offers a new approach *in vitro* to observe the working mechanism of IPC device, offering important data for its further clinical standard regulation.

## Introduction

Deep vein thrombosis (DVT) is the abnormal blood clots formed in the deep veins, often occurring in the perioperative period of major orthopedic surgery^1^. DVT will lead to pain, edema and ulcers, and even disability and death^2, 3^. It has been known that DVT resulted from a combination of flow stasis at the cusps, hypoxia-induced activation of the endothelium and subsequent accumulation of procoagulant facors^4^. In clinical practice, mechanical and pharmacological measures can be used to prevent DVT^5, 6^. Mechanical methods involve the use of intermittent pneumatic compression (IPC), graduated compression stockings (GCS) and venous foot pumps to improve blood circulation^7, 8^. The pharmacological prophylaxis includes aspirin, rivaroxaban, and low molecular weight heparin (LMWH) ^9-11^. Adding these nonsteroidal antiinflammatory drugs (NSAIDs) in patients can reduce the vascular disease but increases the risk of major bleeding^12, 13^. LMWH has been shown a modest reduction in the risk of DVT compared to untreated groups^14^. The poor efficiency is induced by the reduced blood flow to the limb due to immobilisation of the muscle mump, which plays a leading role in DVT pathogenesis. Mechanical measures are used to improve the mulscle pumps, thus increasing the volume flowrate and velocity of blood relux in the vein. Meanwhile, these methods have no risk of bleeding or burden on rivers and kidneys especially for patients at high risk of bleeding complications^15, 16^.

In orthopaedic procedures such as joint arthroplasty, fat droplets, cell clumps and other emboli in the bone marrow cavity can enter the cardiovascular system, increasing the risk of endothelial cell damage and levels of inflammation. The risk of postoperative DVT would be further increased by postoperative bed rWest and limb braking. IPC devices have been widely investigated in human limb and joint treatment since 1960s ^17, 18^. Chibbaro et al. ^19^ divided the patients into two groups A (No IPC), and B (with IPC device). About 3.2% of people in group A suffered from DVT, 0.9% from pulmonary embolism (PE), and 9 people died. In comparison, the number of patients in group B suffered from DVT, PE or death was 0.8%, 0.19%, and 9 respectively. Dennis et al. ^20^ also proved that the application of IPC devices could greatly prevent thrombus for patients with different conditions. Galyaev et al.^21^ confirmed that IPC devices could reduce 62% in the risk of DVT. Nandwana et al.^22^ compared the sequential and single-compartment compressions modes of IPC. The experimental results showed that IPC can increases oxygenation of the peripheral limb muscles, especially during the sequential compression mode. Lee et al. ^23^ introduced a new IPC method, which may improve its safety by maintaining blood flow stability. The compressive force driven by IPC devies defomed the veins, thus in trun cause desirable physiologic effect on preventing thrombus formation^24-27^. However, current studies are still limited to clinical trials. Meanwhile, the standard of pressure and working time intervals of IPC pumps has not been established yet. Thus the use of IPC suffers from lack of consistency in clinical practice.

In addition, the process and mechanism of the IPC pump in preventing lower limb DVT remains unexplored. The emergence of high-precision techniques such as super microvascular imaging (SMI)^28^and Color Doppler^29^ has provided new rearch techniques. However, it is still difficult to precisely observe flow patterns in the venous vein, trajectory of blood cells or the thrombus growth situation in real-time. Furthermore, various flow patterns would be induced in the vein according to different compressive pressure and working time intervals. Microfluidic chip offers a new approach to predict the trends of blood flow^30, 31^. This flexible channel embedded with symmetrical valves can mimick the altered blood flow under compresison and observe the process of thrombis using high speed optical visulaition equipments.

Here, we introduced a microfluidic vein chip to represent the process of thrombosis forming. Finite elenment analyzing modal of the venous vein was established and Computational Fluid Dynamics (CFD) was used to investige the blood behaviour through the vein. The vein chip was then filled with whole blood flow to recapitulate the microenvironment of the actual veins. Further, we determin how the working paramaters of IPC devices alter the blood flow pattern and thus affect DVT formation. This study integrated simulated modals and experimental validation to provide a guidance for clinical IPC usement regulation and innovation of engineered new pneumatic compression equipments.

## Materials and Methods

### Clinical data acquisition

This study was approved by the institutional ethics committee and consents were obtained from all patients. The exclusion criteria were: (1) Patients with fractures, coagulation disorders or known lower limb VTE prior to admission. (2) Incomplete information for any other reason. From September 2013 to August 2015, a total of 543 patients undergoing arthroplasty were recorded in the clinical data, and 278 patients were excluded due to the exclusion criteria mentioned above. In the end, 264 patients were included. From September 2013 to July 2014, patients who used the pneumatic pump for 1 hour per day until discharge were included in the control group. From February 2015 to August 2015, patients who used the pneumatic pump for 24 hours on the first postoperative day and then for 8 hours per day until discharge were included in the IPC group.

### Finite element analysis

Doppler Ultrasound image scans were performed with patients in a relaxed supine position and the blood flow velocity was measured at a point between the venous valves of the femoral vein. The mass and momentum equations were solved by the CFD software COMSOL Multiphysics (v5.5) in 2D based on finite element analysis. The geometric modal of the vein was established consisted of two pairs of valves and a long microfluidic channel originating from patients’ veins. The valve leaflets were considered as linear elastic materials with Young’s Modulus of 100kPa^32, 33^. Blood was considered as a non-Newtonian fluid with constant density of 1050 kg/m^3^ and Reynolds number of 1.05 closing to numan blood. The blood flow was considered to be an incompressible fluid flow. The vein were squeezed by IPC device periodically including the time for one working cycle and the time interval between two working cycles. During the working cyle, the inlet boundary condition was fitted with piecewise funtions based on the clinical measured data. At the interval period, the blood would flow smoothly in the vein channel. The valve leaflets were constrained to the vein wall.

Poiseuille’s law was applied to describe the laminar pressure confined to vein channel. The blood flow rate was generated by pressure differences at two points of the vein. The relationship between the blood flow velocity *v* and the radius *r* can be given as^34^:

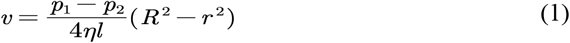

where *l, p*_1_*-p*_2_, *η*and *R* respectively represented the the length of the vein channel, the pressure gradients between the two ends of the vein channel, and the inner radius of the vein channel.

### Fabrication of microfluidic vein chip

The model of the vein chip was created by CAD software (Autodesk). The body of the vein chip was made by etching channels on the PDMS using soft lithography technology. These PDMS blocks utilized three parallel channels per chip, each having an inlet and outlet hole with a 1.0mm width. The opening of each channel was about 50% of the width of the channel. Glass glue was used to bond PDMS blocks and slides together. Vein-chips with glass or not were used for blood perfusion experiments and VWF expression respectively.

### Blood sample collection, treatment and testing

5mL of blood was collected from the patients’s median elbow vein the day after total joint arthroplasty and stored in a vacuum tube containing 3.2% sodium citrate to prevent coagulation. Informed consent was obtained from all donors for the collection and use of all blood samples. FITC-labeled human fibrinogen (final concentration 12.5μg/ml) was added to the collected blood to visualize the deposition of the fibrin.

Platelets were labeled with the FITC labeled CD41 Monoclonal Antibody (final concentration 5μg/ml). The labeled blood was incubated at 37°C for 15 minutes prior to the assay to give the FITC a clearer fluorescence under blue light excitation. The labeled blood sample was prepared by rapid mixing with recalcification buffer (75 mM CaCl_2_ in PBS) to restore the coagulation. The channel was perfused with a PBS solution containing 2% bovine serum albumin (BSA) before infusing blood into the vein chip. The process of venous thrombosis was measured using fluorescence microscopy (IX83, Olympus Co.)

The venous blood was centrifuged at 3000rpm for 15min. The centrifuged RBCs were put into the PBS solution with a hematocrit value of 0.5%. Under an inversion microscope, RBCs were used directly as tracer particles to illustrate the flow field in a bright field. The movement of the RBCs under the gradient pressure of the controller (Elvesys Co.) was recorded by a high-speed camera (M110, Phantom Co.). The recorded videos were analyzed to acquire the velocity field of the fluid (Fig S1). The capture speed of the high-speed camera was 2000fps. The blood flow velocities in the veins at different IPC pressures were obtained based on preliminary clinical data. When observing venous thrombosis, the parameters of the console were adjusted so that the blood flow velocity in the microfluidic channel corresponded to the venous blood flow velocity.

### Cell culture and immunohistochemistry

A microfluidic pump was used to infuse the dopa solution (2mg/mL in the 10 mM Tris-HCL, pH 8.5) into the microfluidic channel at 10μL/min until the channel was filled with dopa solution. After standing for 5min, the excess dopa solution was flushed out of the tubing using PBS. The gelatin from porcine skin was dissolved in PBS to make a 10% solution and instilled into the channel. Keeping for 15 minutes, the channels were flushed with PBS for 5min. The encapsulated chips were irradiated under UV light for 2 hours and used for cell culture.

Human umbilical vein endothelial cells (HUVECs) were cultured in DMEM medium containing 10% v/v fetal bovine serum (FBS) and 1% v/v penicillin-streptomycin solution at 37? under 5% CO_2_. The cells were digested down to prepare a suspension when they reached 90% confluence. The HUVECS suspension (2×10^7^/mL) was infused into the chip channels at a rate of 2uL/min. The chips were placed in a cell incubator for 48 hours and fresh medium was injected into the tube every 12 hours using a microfluidic pump. Excess medium was kept at the inlet and outlet of the chip channel to prevent evaporation of liquid from the channel. After 48 hours, HUVECs were observed adhering and growing inside the chip channel and forming good cell junctions. A microfluidic pump was used to instill the medium containing 5ng/mL TNF-α into the channels at different rates to simulate the effects of different blood flow rates on venous endothelial cells.

Vein-chips were fixed with 4% paraformaldehyde solution for 30 minutes. Then the vein-chips were washed with PBS 3 times (5 minutes each time). 0.5% Triton X-100 was gently added dropwise to the chip surface and evenly covers the cells for 15 min. 5% BSA (based on PBS, room temperature for 60min) with Triton was gently dripped and evenly covered on the chip surface. The 1:200 Anti-Von Willebrand Factor antibody (ab6994, Abcam) was configured with 5% BSA and uniformly covered on the chip surface and incubated at 4°C for 24 h. After washing, the 1:400 goat anti-rabbit IgG H&L (ab150077, Abcam) of PBS configuration was uniformly covered on the surface of the chip channel and incubated for 120 minutes at room temperature. Then it was washed with PBST and covered with DAPI.

### Blood flow velocity measurement

With the inverted microscope (CKX41, Olympus Co.), we regarded the width of the channel as the standard length to measure the velocity of RBCs in a bright field. The process of flowing RBCs was recorded by a high-speed camera (M220, Qian Yanlang Co.) with 2000 fps. It was hard to synchronously measure the velocity of blood when it flowed. All the measuring data were recorded for many times and reached an average number then. The experiments were operated with four models: patients only wearing elastic stockings and with IPC under pressure of 20mmHg, 40mmHg, and 60mmHg. The cycle period was separately set up as 7.2s?8.2s and 9.8s corresponding to 20mmHg, 40mmHg and 60mmHg. Finally, we chose the distinctive RBCs as points of speed measurement and average the speed of RBCs.

## Results

### Clinic Data

In this study, a total of 264 patients were enrolled. 135 patients were included in the control group and used the pneumatic pump for 1 hour per day until discharge. 129 patients were included in the pump group. They used the pneumatic pump for 24 hours on the first postoperative day and then 8 hours per day until discharge. The mean age of the control group and the IPC group were respectively 63.79±11.49 and 64.74±10.55 years old. The average D-dimer of the control group and the IPC group before surgery were respectively 0.95±1.16 and 1.05±1.76 (Fig S2). There was no statistical difference between these two groups (P>0.05). Detailed data were listed in Table 1.

**TABLE 1.** Basic clinical characteristics of patients involved in this study

| Variables | Control<br>(Mean ± SD) | IPC<br>(Mean ± SD) | P value |
| --- | --- | --- | --- |
| Age (years) | $63.78 \pm 11.49$ | $64.74 \pm 10.55$ | P=0.485 |
| BMI (kg/m<sup>2</sup>) | $25.14 \pm 4.01$ | $25.81 \pm 3.87$ | P=0.173 |
| Gender (female) | 90 (66.7%) | 96 (74.4%) | P=0.169 |
| Fbg (g/L) | $3.00 \pm 0.60$ | $2.97 \pm 0.73$ | P=0.378 |
| D-dimer (mg/L) | $0.95 \pm 1.16$ | $1.05 \pm 1.76$ | P=0.468 |

A total of 27 patients (20.0%) in the control group were found to have deep venous thrombosis (DVT) by venography and ultrasound findings. Soleal vein (SV) thrombosis occurred in 22 patients (16.3%), peroneal vein (PEV) thrombosis in four patients (3.0%), posterior tibial vein (PTV) thrombosis in two patients (1.5%) and femoral vein (FV) thrombosis in one patient (0.7%). One patient had a combination of PTV thrombosis, PEV thrombosis and SV thrombosis. In the IPC group, 19 patients (14.7%) were identified as having DVT, including 18 cases (14.6%) of SV thrombosis, and one PTV thrombosis (0.8%). There was no significant difference in the incidence of deep vein thrombosis between the two groups (P=0.326). However, the incidence of major venous thrombosis (PEV+PTV+FV) was significantly lower in the IPC group than in the control group (P<0.05). The specific distribution of thrombosis was shown in Table 2. On the day of venography, plasma D-dimer levels were 3.34±1.94 (mg/L) in control patients and 2.53±1.68 mg/L in the IPC group. It is well known that high D-dimer levels associate with risk factors for thrombosis. It indicated that the risk of thrombosis can be reduced using IPC, which was consistent with the results of venography. The comparison between this study and other similar studies in which IPC reduced DVT after arthroplasty was listed in Table 3.

**TABLE 2.** Distribution of different kinds of DVT among the patients

| Variables | Control<br>N (%) | IPC<br>N (%) | P value |
| --- | --- | --- | --- |
| DVT | 27 (20.0%) | 19 (14.7%) | 0.326 |
| SV | 22 (16.3%) | 18 (13.9%) | 0.596 |
| PEV | 4 (3.0%) | 0 (0%) |  |
| PTV | 2 (1.5%) | 1 (0.8%) |  |
| FV | 1 (0.7%) | 0 (0%) |  |
| PEV+PTV+FV | 7 (5.2%) | 1 (0.8%) | 0.048 |

**TABLE 3.** Comparison between this study and other similar studies in which IPC reduced DVT after arthroplasty. (N/A=not available)

|  | **This study** | **Kim, 2019** | **Liu, 2017** | **Jo, 2016** | **Tsuda, 2015** |
| --- | --- | --- | --- | --- | --- |
| Surgery type | THA+TKA | TKA | TKA | THA | THA |
| IPC application pressure | 40mmHg | N/A | 45 mmHg | 30-45mmHg | 110mmHg |
| IPC application time | 5 days | 6 days | 48 hours | 2 days | 1-2 days |
| IPC application site | Calf | Ankle to thigh | Calf and thigh | Ankle to thigh | Foot |
| Additional measures | Anticoagulants | Compression stockings | Anticoagulants | Anti-embolic stockings | N/A |
| DVT detection after operation | 5 days | 6 days | 9 days | 5 days | 3, 21 days |
| Number of participants | 129 | 425 | 60 | 741 | 166 |
| Incidence of overall DVT (%) | 14.7 | 11.3 | 8.3 | 0.4 | 6.0 |
| Proximal DVT (%) | 0 | 0.9 | 0 | N/A | 0 |
| Distal DVT (%) | 0.8 | N/A | 5 | N/A | 6 |
| Intermuscular DVT (%) | 13.9 | N/A | 13.3% | N/A | N/A |

### Design and Fabrication of Microfluidic Vein Chip

We firstly needed to acquire the anatomy of a human vein and decide the geometric parameters of a microfluidic vein chip. Thus, we analyzed the Doppler-ultrasound images at the femoral vein with Portable Color Doppler (PCD, M-turbo, SONOSITE Co.) (Fig 1A). The vein valve consisted of two leaflets which opened when the blood flowed across them. Considering the maximum width of the venous valve, the distance between the valve leaf tip and the valve bulb, the channel dimensions were as follow (Fig 1B). The width *L* and depth of the channel were selected to be 350μm and 50μm respectively. The gap width between the leaflets *d* were designed to be 160μm. The width of the venous valve bulb was set at 600μm. Then we fabricated the vein chip. Three channels with a length of 40mm and separated by 5mm were placed in a vein chip device (Fig 1C and Fig S3). Figure 1D showed the meshed domain using free triangular meshes.

**FIGURE 1.**
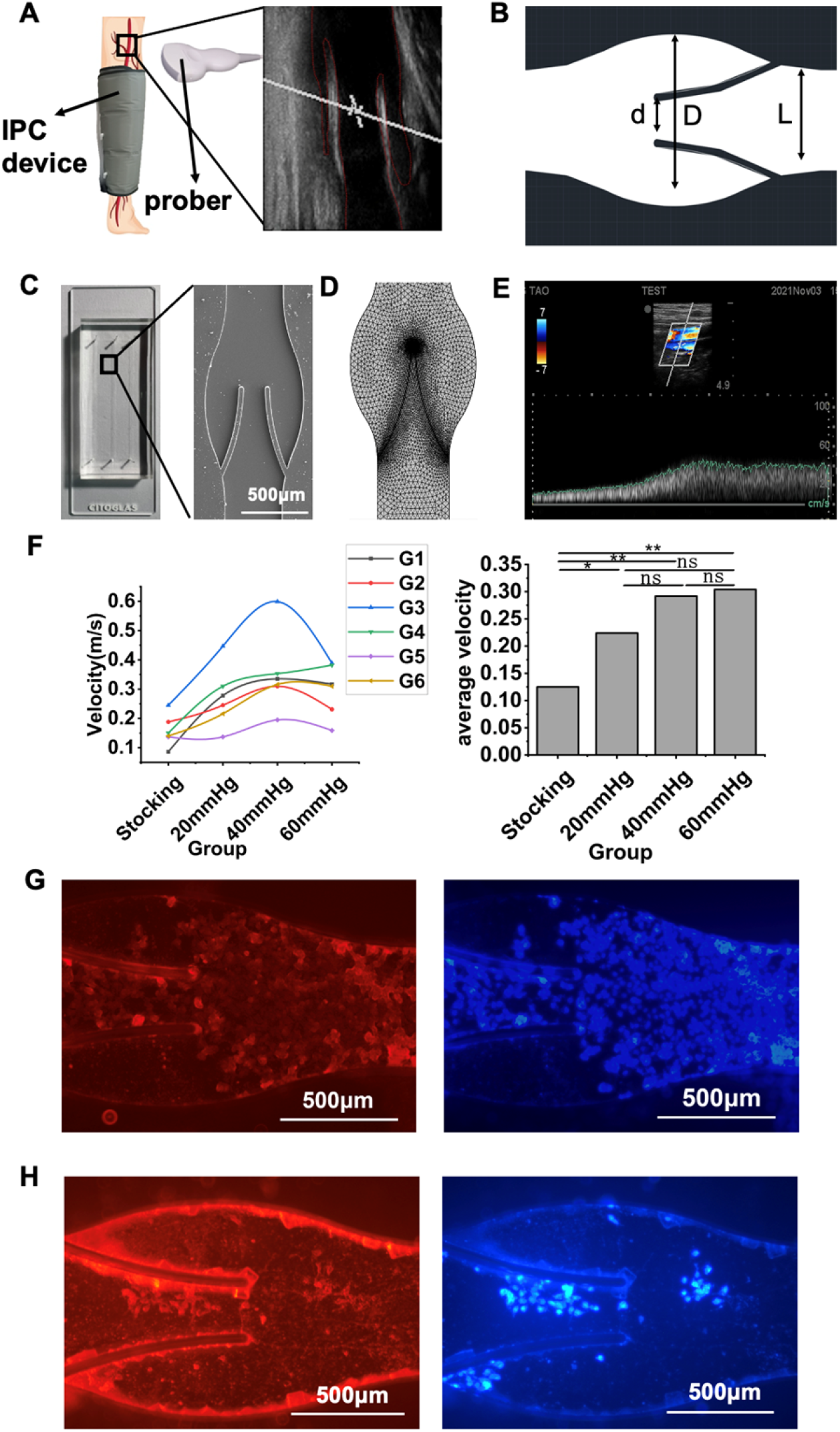
Design and analysis of the vein chip. **A** Illustration of working principles of IPC treatment and Doppler-ultrasound image of the vein. **B** CAD figure of the vein chip. **C** Photograph and Electron microscope view of the vein chip. **D** Mesh elements of the vein channel. **E** Velocity between leaflets obtainded by Doppler-ultrasound image. **F** Maximum and average blood flow rate among the patients with or without IPC. Fluorescence micrographs showing immunostaining of von Willebrand Factor (VWF, red) and DAPI staining (blue) of the venous valve with entrance flow rate simulating **G** IPC. **H** control group.

Color Doppler ultrasound was performed on the femoral vein on the surgical side and healthy side respectively (Fig S4). The supine position was taken during operation to prevent the effect of posture on femoral flow velocity, ensuring that the patient can breathe steadily without talking. Then the velocity of the vein was measured. We monitored patients of different ages and genders after knee or hip replacement. No obvious difference of blood flow velocity of the femoral vein was observed between the surgical side and healthy side. In Fig 1E, G1 referred to a male thin patient, G2 and G3 represented the healthy side and the perioperative side of another slim man respectively, G4 and G5 referred to the healthy side and the perioperative side of another fat woman, G6 was a moderate-sized woman. As shown in Fig 1E and Table S1, the maximum and average blood flow velocity were both higher with IPC with 40 mmHg or 60mmHg maximum pressure value. The results also showed that patients with slow basal flow rates may produce a higher peak flow rate with IPC.

VWF is considered an initiator of platelet adhesion and aggregation, leading to thrombus formation. Thus, we measured quantities of VWF in the vein chip. We observed that the expression of VWF was less in the IPC group compared to the control group (Fig 1G). To further characterize the cell adaption to the micro-channel in vein-chips, 4’,6-diamidino-2-phenylindole (DAPI) was used to survey the living cells (Fig 1H). The results further confirmed that the high share and flow rate caused by IPC compression could reduce platelet adhesion and thrombus formation.

### CFD Analysis of Vein Chip

According to different treatments with IPC, we divided the clinical patients into 4 groups: NIPC (no treatment with IPC), 20mmHg, 40mmHg and 60mmHg (the maximum compressive pressure of IPC). Patients in both NIPC and IPC groups worn the elastic stocking on the lower limb in the experiment. CFD techniques were conducted on the modal of the femoral vein. The blood flow rate was modeled using mathematical functions mimicking the clinical situation. The maximum velocities for the IPC group (20mmHg, 40mmHg or 60mmHg) were respectively 1.6, 1.7, and 1.8 times compared to that of the NIPC group. Then the shear stress and laminar pressure in the vein were studied.

The blood flow behavior aound the valve and the flow patterin were shown in Fig 2A. Through the main channel, a high-speed jet was formed. At the same time, two contra rotating vortex can be seen at the venous valve cusp. Fig 2B showed the shear stresse around the venou valve leaflet when the valve was fully opened. The maximum shear stress happened at the connection point of the valve to the wall reached largest, and then decreaed till the tip of the valve leaflet. The reflux can reach the bottom of the pocket and increase the flow range in this region. The flow pattern as the leaflets opened versus time was shown in Fig 2C. In the beginning, the valve was closed and no blood flew in the channel. As inlet velocity increased, the blood velocity rised from the inlet passing the valve and then flew toward the outlet. The valve opening increase as a result of the increasing velocity. The velocity reached a maximum magnitude where the cross-sectional area between the vein channel walls was minimum. It can be seen in Fig 1E that the maximum velocity happened corresponding to 89.5% of valve opening. After that, the valve opening decreased by the time.

**FIGURE 2.**
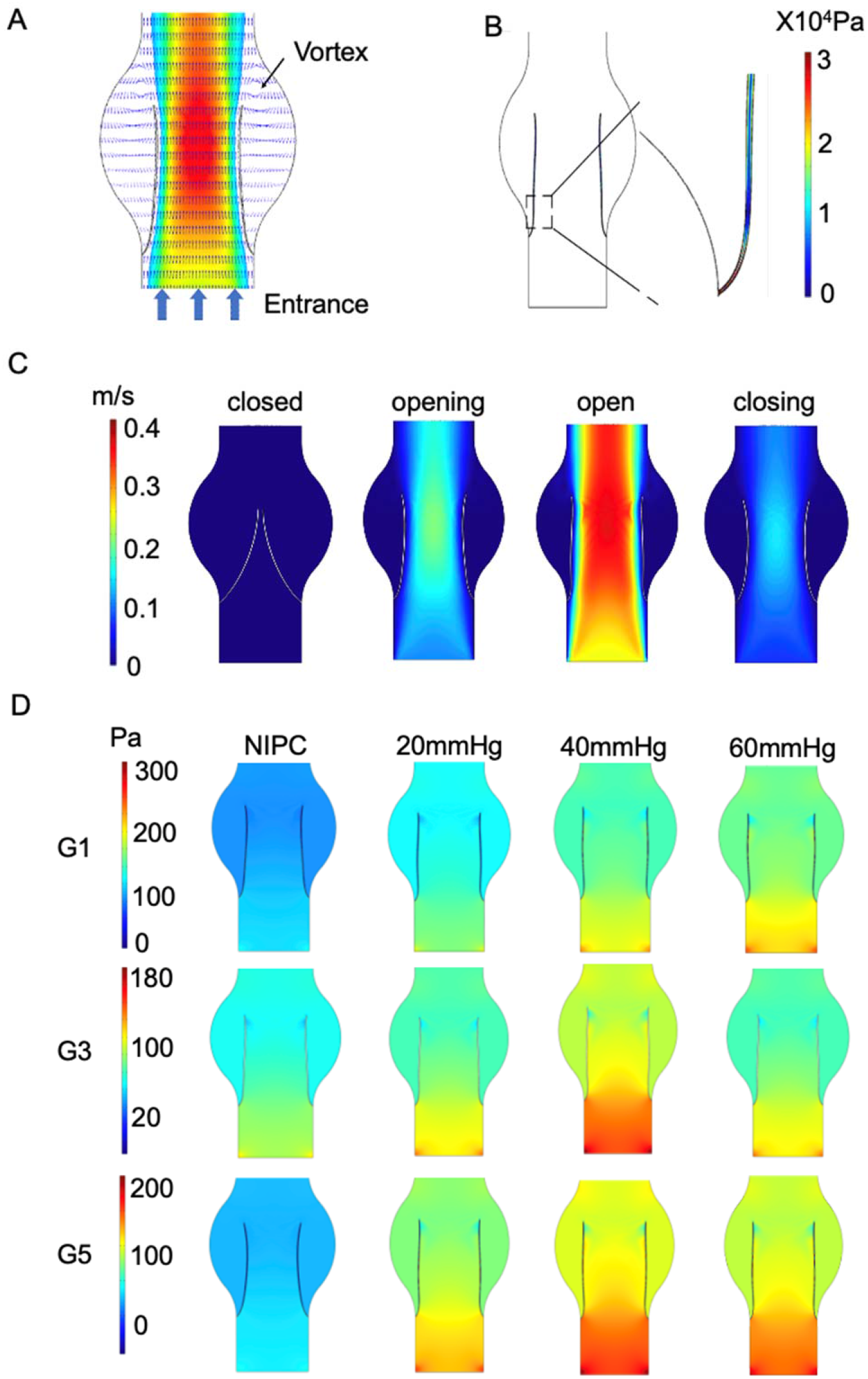
Contour maps showing pressure and velocity from CFD results. **A** Velocity contours around the valve. **B** The shear stress on one side of the venous valves. **C** Velocity variation when leaflets open through time. **D** Laminar flow pressure within the vein channel.

Laminar low is determined by concentric layers of blood moving inparallel along the vein. Fig 2D showed the steady flow conditions of the patient G1, G3 and G5 when the valve was fully opened as well. The maximum laminar flow pressure was found in the center of the vein channel while the lowest velocity was found along the channel wall. The distribution of the laminar flow pressure was in accordance with that of the flow velocity. Parabolic flow was hardly seen as for the control group. On the contrary, steay laminar flow can be found among all the IPC groups. The blood flow under 40mmHg showed maximum value of laminar flow pressure. No obvious difference was seen for patient G3 with IPC treatment under 20mmHg and 60mmHg.

### Thrombus growth under different operating modes of IPC

Platelets and fibrins are commonly used markers to visualize thrombus formation. The growth of the thrombus in the valve pocket was investigated by observing the deposition of fibrinogen and platelets using fluorescent microscopy. The growth of the thrombus in 20 minutes in the valve pocket without or with different maximum IPC pressure levels were analyzed. The IPC device worked with a constant time interval of 48s. As shown in Fig 3A, red blood cells (RBCs), platelet and fibrin began to accumulate at the time of 20 minutes, especially in the valve pocket. The number of fibrin in the vortex zone was significantly lower in all of the IPC group than in the control group (p<0.01). The fibrin accumulation could be lessened with the increase of IPC compressing pressure (Fig 3B and Fig 3C). The platelet adhesion within the valve pocket also decreased due to higher IPC working pressure (Fig 3D and Fig 3E). It can be concluded that the altered blood flow driven by IPC took effect on the thrombus formation in venous valves. Both of the blood flow rate and the shear stress became higher as the IPC pressure increased. Although more platelets might be activated due to higher shear stress^30^, the fibrin monomers and oligomers became unstable, still reducing the thrombus deposition. Meanwhile, the stagnation of blood flow might lead to release of inflammatory factors, which could also increase the thrombosis deposition.

**FIGURE 3.**
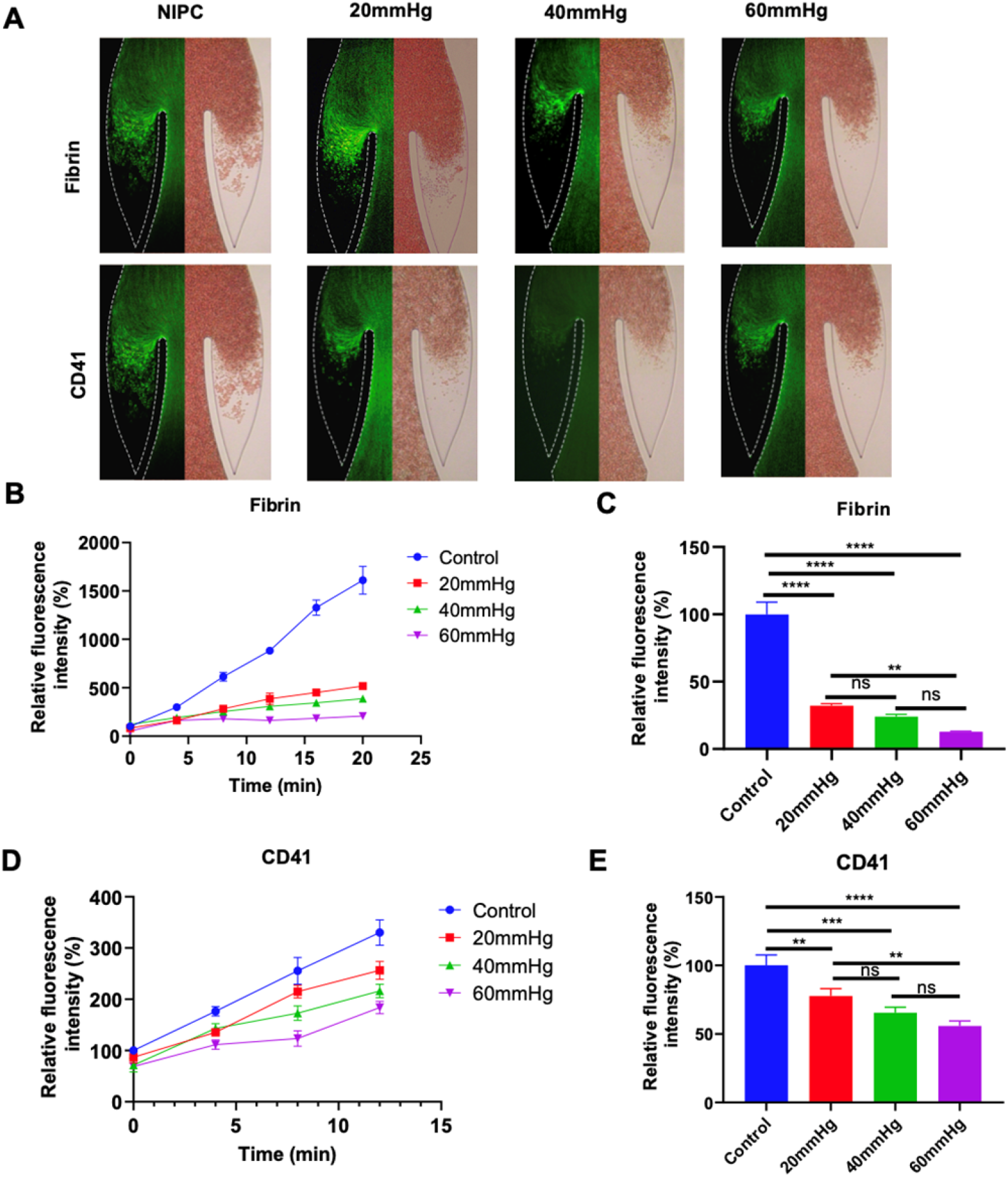
Thrombus deposition progress in altered blood flow under different IPC working pressure. **A** Fluorescence micrographs of Vein Chips after perfusion of blood labelled with fibrin (top), platelets (bottom) for control and different IPC groups. The relative intensity of fibrin. **B** over 20 minutes, and **C** at 12^nd^ minute. Relative intensity of platelet **D** over 20 minutes, and **E** at 12^nd^ minute.

The growth of thrombus in 20 minutes in the valve pocket with IPC treatment under different time intervals was analyzed as well (Fig 4A). The IPC device works with constant working time 12s and different interval times including 24s, 48s and 60s. At the initial time, there was a small are of thrombus found in the valve pocket. The thrombus grow with the increase of interval time(Fig 3B-3E). The fibrin and the platelet accumulated in the valve at the 12nd minute were almost twice with the interval time 60s as those with the interval time 24s (Fig 4C and Fig 4E). No obvious difference was found between the 12+24 group and the 12+48 group in terms of reducing fibrinogen deposition (p>0.05). In contrast, a significant difference between the 12+24 group and the 12+48 group in terms of reducing platelet deposition in the valve area (p<0.01). The results indicated that the altered blood flow induced by longer interval time of IPC was more important to platelet accumulation than fibrin. IPC device with the working time of 12s and interval time of 24s was considered as the most optimal parameters for preventing thrombus deposition.

**FIGURE 4.**
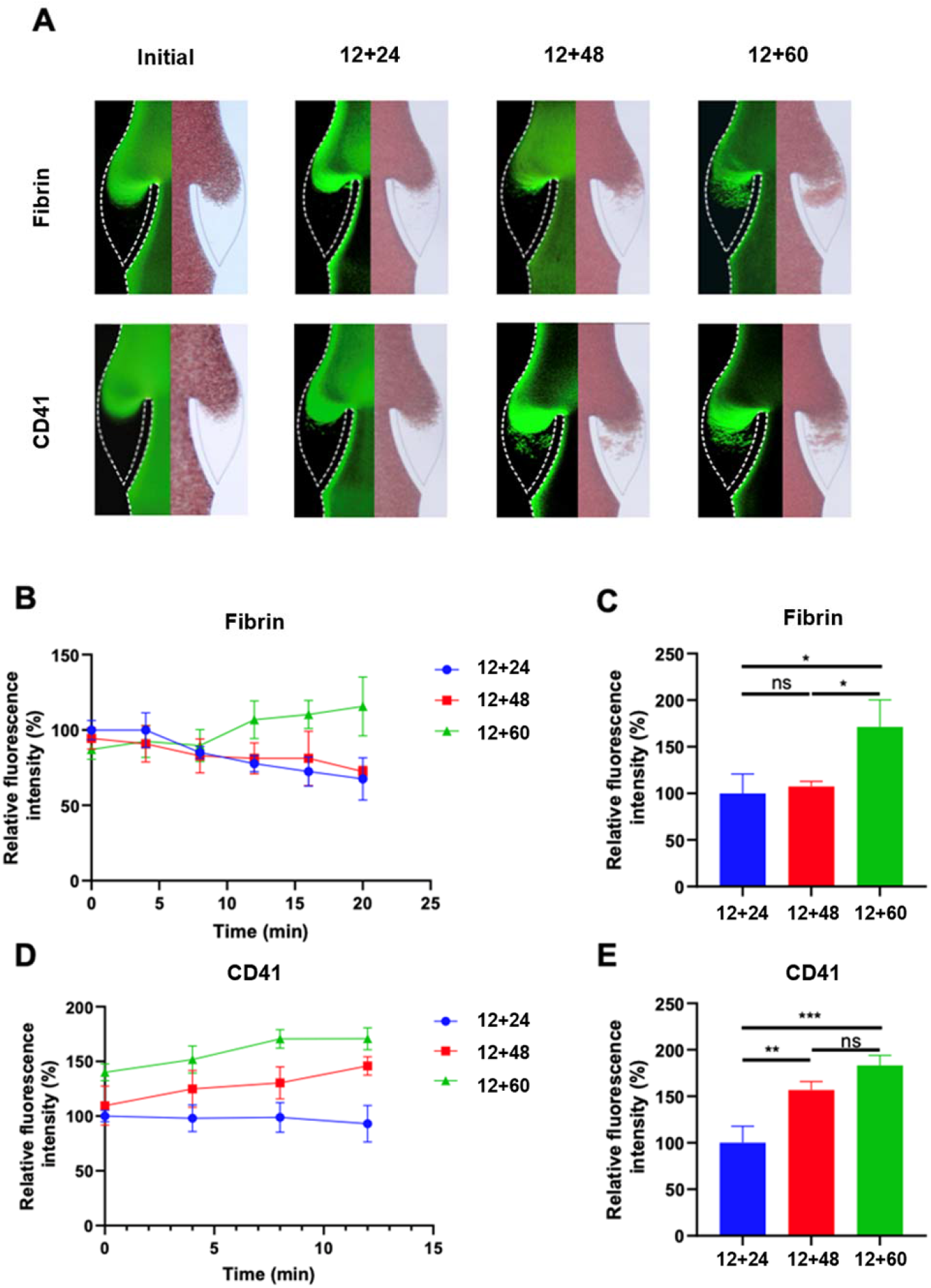
Thrombus deposition progress in altered blood flow under different IPC time intervals. A Fluorescence micrographs of Vein Chips after perfusion of blood labeled with fibrin (top), platelets (bottom) for control and different IPC groups. Relative intensity of fibrin B over 20 minutes, and C at 12^nd^ minute. Relative intensity of platelet D over 20 minutes, and (E) at 12^nd^ minute.

## Discussion

DVT often happens in the lower limb veins in patients restricted to bed after hip or knee replacement surgery^35^. Trunk vein (PEV, PTV, FV) thrombosis might lead to severe disease such as pulmonary embolism. We have conducted hundreds of analyses on thrombus formation of patients with IPC treatment after knee surgery from September 2013 to August 2015. This prophylactic measure, mimicking the natural muscle pumps, had been proved to reduce the incidence of DVT. However, the working mechanism how IPC influence the blood flow in the veins of the lower limbs has rarely been researched. In this work, we constructed a microfluidic chip system based on a simulated human lower limb vein to observe the process of venous thrombosis in real time and to assess the differences in thrombosis under different IPC parameters. The microchip has a similar structure to the valve region of the lower limb deep veins and can simulate the flow in the valve region where thrombosis most commonly occurs. In addition, the computer simulations as well as the blood flow rates in the microfluidic channels of the chip were derived from Doppler ultrasound measurements of venous blood flow rates in the subject’s lower limbs. Also, blood samples from patients after surgery were used to mimic the inflammatory conditions that the blood and endothelial cells experience. By combining data simulation with actual measurements, the microchip system is expected to simulate venous thrombosis from the perspectives of valve structure, flow velocity, and the inflammatory environment. It will also look into the impact of various IPC modes on the intervention of venous thrombosis in the lower limbs.

The compressive pressure amplitude and the frequency of the IPC device were researched in this study. The maximum pressure ranged from 20mmHg to 60mmHg. The working time was set as 12s for all the IPC groups, while the time intervals varied including 24s, 48s and 60s according to the commercial device. Immobility can lead to low-speed blood flow and limitation of blood exchange within valve pocket. It was noticed that the blood stasis in the valve pocket tended to form without IPC compression. When the IPC works periodically, vortex in the valve pocket was generated. The velocity reached to its maximum value passing the valve and decreased along the walls. It was led by the decrease of flow cross-section area and vice versa. Although vortices can be seen behind the leaflets, their velocity were lower, especially in the region close to the connection point of the valve to the vein wall. The vortices became more obvious towards the valves tips.

Separated clots can be moved by the blood flow. With larger compressive pressure or shorter time intervals between the working cycles, the increased flow velocity could reduce the fibrin and platelet deposition in the valve. Without enough shear stress and flow veloticy, blood stasis would lead to thrombus formation. As the compressive pressure declined or the time intervals increased, the vortex could not destroy the network composed of cross-linked fibrin, aggregated platelets and red blood cells^36^. The results showed that high blood flow pressure restrained more fibrin linkage while short time interval controlled recruitment of platelets. By including venous valves and blood flow in this vein chip, we observed optimal working parameters for IPC devices. This data may be useful in designing future therapertic methods.

However, the vein chip platform still has limitations that need to be take into consideration in future work. The blood samples used in the experiment were stored in the sodium citrate to anticoagulation and recalcified when used. The excessive calcium might lead to changes in platelet function^37^. We also noticed that thrombus formed faster in the microengineered vein chip than in the actual vein in clinical. It is possibly caused by the lack of the other two factors in thrombus formation: endothelial dysfunction and blood hypercoagulations. The endothelial cells within the vein significantly regulate blood cell adhesion, vascular inflammation and thrombosis formation^38^. Hypoxia conditions in the chip channel may also induce the release of the inflammatory mediators that would increase the thrombus formation^39^. Meanwhile, the valve leaflets in this chip were fixed due to limited manufacturing technology^40-43^. The natural venous valves are flexible instead, allowing more blood flowing through the vein when they are opened. The mechanical modulus, the gap distance and the shape of the valve leaflets also impact the cell or protein attachment^32^.

## Conclusions

In summary, this microfluidic vein chip firstly allowed observation of IPC working mechanism on preventing DVT *in vitro*. Using this device, we could illustrate how the vein valves behave under IPC compression and how the altered blood flow influences the thrombus formation in the vein. The obtained data would be helpful in regulating clinical operation standard of IPC devices in the future.

## Supplementary Information

Additional file 1. Supplementary Material.

**FIGURE S1** Captured images of red blood cells for velocity measurement. **FIGURE S2** D-dimer levels of blood collected from the control and IPC group patients. **FIGURE S3** CAD drawings of vein chips integrated three channels. **FIGURE S4** Doppler ultrasound images of the femoral vein. **A** Patients only wearing the elastic stockings. Patients taking IPC treatment with the maximum pressure of **B** 20mmHg **C** 40mmHg and **D** 60mmHg. **FIGURE S5** Simulation of laminar flow pressure within the vein channel of patient G2, G4 and G6. **TABLE S1** Peak blood flow rate for IPC group.

## Declarations

### Ethics approval and consent to participate

All experiments were conducted in accordance with the guidelines approved by the Ethics Review Committee of Nanjing Drum Tower Hospital.

### Consent for publication

Yes

### Availability of data and materials

The datasets used and/or analyzed during the current study are available from the corresponding author on reasonable request.

### Competing interests

The authors declare that they have no competing financial interests or personal relationships that could influence the work reported in this paper.

### Funding

This work was supported by the National Natural Science Foundation of China (32171358, 52005268), Graduate Scientific Research and Innovation Plan Project in Jiangsu Province (KYCX22_1621), the Natural Science Fund for Colleges and Universities in Jiangsu Province (20KJA460004), the Open Research Fund of Guangdong Key Laboratory of Minimally Invasive Surgical Instruments and Manufacturing Technology, Guangdong University of Technology (MISIMT-2021-5).

## Authors’ contributions

H.D, Y.Y and L.Z designed the study. H.D and S.C performed the experiments. J.S and W.T collected the data. S.C analyzed the data. H.D and L.Z wrote the manuscript. Y.Y and J.Y revised the manuscript. Q.J and L.Z supervised the study.

## Acknowledgements

None

## Notes

### Competing Interest Statement

The authors have declared no competing interest.

